# Endocytosis gated by emergent properties of membrane-clathrin interactions

**DOI:** 10.1101/2023.08.02.551737

**Authors:** Xinxin Wang, Yueping Li, Ailing Liu, Ronnin Padilla, Donghoon M. Lee, Daehwan Kim, Marcel Mettlen, Zhiming Chen, Sandra L. Schmid, Gaudenz Danuser

## Abstract

Clathrin-mediated endocytosis (CME), the major cellular entry pathway, starts when clathrin assembles on the plasma membrane into clathrin-coated pits (CCPs). Two populations of CCPs are detected within the same cell: ‘productive’ CCPs that invaginate and pinch off, forming clathrin-coated vesicles (CCVs) [1, 2], and ‘abortive’ CCPs [3, 4, 5] that prematurely disassemble. The mechanisms of gating between these two populations and their relations to the functions of dozens of early-acting endocytic accessory proteins (EAPs) [5, 6, 7, 8, 9] have remained elusive. Here, we use experimentally-guided modeling to integrate the clathrin machinery and membrane mechanics in a single dynamical system. We show that the split between the two populations is an emergent property of this system, in which a switch between an *Open* state and a *Closed* state follows from the competition between the chemical energy of the clathrin basket and the mechanical energy of membrane bending. *In silico* experiments revealed an abrupt transition between the two states that acutely depends on the strength of the clathrin basket. This critical strength is lowered by membrane-bending EAPs [10, 11, 12]. Thus, CME is poised to be shifted between abortive and productive events by small changes in membrane curvature and/or coat stability. This model clarifies the workings of a putative endocytic checkpoint whose existence was previously proposed based on statistical analyses of the lifetime distributions of CCPs [4, 13]. Overall, a mechanistic framework is established to elucidate the diverse and redundant functions of EAPs in regulating CME progression.

## Main

Cells engulf cargo mainly via clathrin-mediated endocytosis (CME), which occurs when nascent clathrin-coated pits (CCPs) evolve into clathrin-coated vesicles (CCVs) [1, 2, 14] (Fig. 1A). Biochemical analyses of CME have identified numerous and diverse endocytic accessory proteins (EAPs) implicated in this progression [6, 7, 15], as well as their interactions [16, 17, 18], and temporal appearances [5, 19]. The process of CME is mechanically demanding as it involves the generation of small vesicles from a flat cell surface. The mechanical demands are met by sufficient chemical energy derived from assembling clathrin coats [20] and association of membrane curvature-generating EAPs [21, 22]. Indeed, a notably high proportion of CCPs, referred to as abortive pits [3], fail to progress towards productive, cargo-loaded CCPs. Based on statistical and kinetic arguments, previous studies have proposed that turnover of abortive CCPs vs. maturation of productive CCPs is gated by a regulatory mechanism, referred to as the endocytic checkpoint [4, 13] (Fig. 1A).

**Figure 1:**
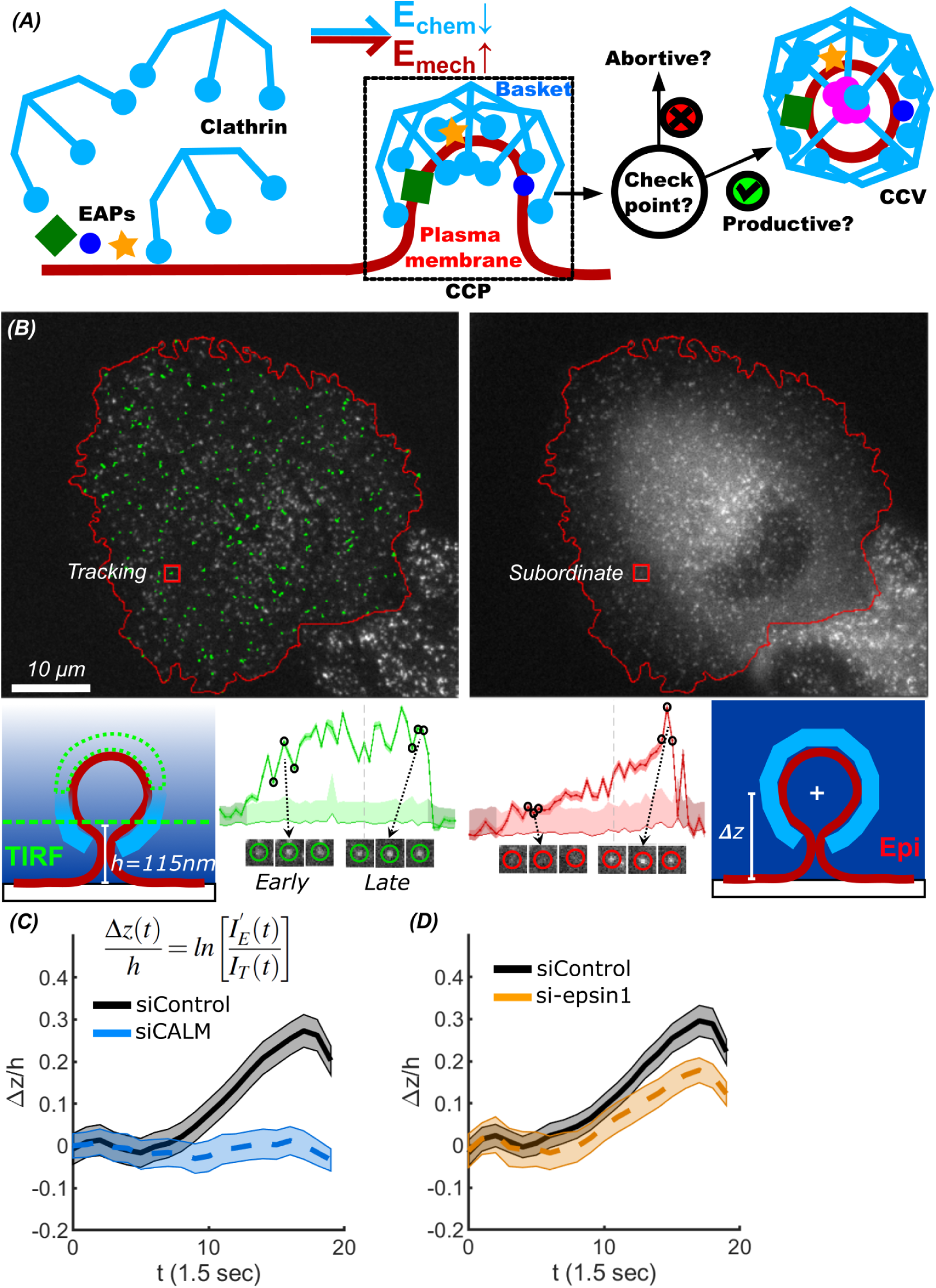
Endocytic accessory proteins (EAPs) initiate curvature generation at clathrin-coated pits (CCPs). (A) Schematic diagram of abortive and productive CCPs and the role of a putative endocytic checkpoint in gating the progression of CCP maturation. (B) Application of Epi-TIRF microscopy measuring curvature formation of CCPs using primary (TIRF-channel)/subordinate (Epi-channel) tracking. Cell border is delineated by red lines. *h* indicates the evanescent depth of TIRF field. ‘+’ indicates the location of the center of mass of the clathrin basket. Sample tracks (green) and a select trajectory (red box) is presented below the images as intensity traces in green (TIRF) and red (Epi). (C-D) Normalized and averaged traces of invagination depth, Δ*z*(*t*)*/h*, measured in (C) siControl (pooled from *n* = 13.7 *×* 10^3^ traces from *m* = 15 cells) and CALM KD (pooled from *n* = 3.1 *×* 10^3^ traces from *m* = 16 cells); and (D) siControl (pooled from *n* = 8.9 *×* 10^3^ traces from *m* = 14 cells) and epsin1 KD (pooled from *n* = 5.6 *×* 10^3^ traces from *m* = 14 cells) conditions. Time is indicated in frames, sampled at 1.5 sec/frame. *I_T_* (*t*) indicates cohort-averaged CCP intensity from TIRF-traces; 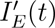 indicates cohort-averaged CCP intensity from Epi-traces subject to a linear correction, see more detail in [28]. Solid/dashed lines indicate the average traces, shaded bands indicate 95% confidence intervals.

The activities of multiple EAPs impinge on the endocytic checkpoint [2, 9, 23]. Yet, in systematic screens, knockdown of most EAPs yield mild or undetectable effects on CME [24, 25, 26, 27], and multiple EAPs have overlapping phenotypes even under sensitive and comprehensive imaging [9]. Indeed, measurements of the effect of EAP knockdown on CME did not correlate with their effects on CCP maturation [28]. Moreover, CME persists in cells expressing a truncated mutant of adaptor protein complex 2 (AP2) deficient in EAP recruitment, yet these cells exhibit dramatically increased numbers of abortive CCPs that fail to gain curvature, despite significant clathrin assembly [13]. These data establish the resilience of CME and suggest the functional redundancy of EAPs. Given this, biochemistry-based approaches alone are insufficient to reveal the regulatory mechanism governing CME progression. Thus, while the recruitment of EAPs and the acquisition of curvature appears to play a role in passing the endocytic checkpoint, the actual gating mechanism has remained elusive.

Other studies have pursued the mechanical aspects of CME by inferring the strengths of clathrin coats on the plasma membrane [29] and purified clathrin baskets [30] from structural studies, as well as from changes in membrane morphology [31], the spatial distributions of EAPs at CCPs [32], and the elasticity of the clathrin baskets [33]. However, these data have yet to be integrated with the known biochemistry to define the endocytic checkpoint and to clarify the conditions of CCP progression. To do so will require quantifying and understanding membrane mechanics, biochemistry, and the interplay between the two in one dynamical system. A data-driven mechanochemically explicit model that includes membrane, clathrin, and EAP components is poised to fill this gap. Existing models [34, 35, 36, 37] lack the explicitness and experimental scrutiny to define the checkpoint from the integrated biochemical and biomechanical properties of CME.

## EAPs initiate curvature generation at CCPs

Abortive pits fail to generate curvature [13, 28]. Additionally, so-called pioneer EAPs that have curvature generating activities [7] have been identified at nascent CCPs [1, 38]. Therefore, to test the role of EAPs in generating early curvature and establishing initial conditions for coat assembly, we measured the time-dependent invagination depth at individual CCPs as an estimation of curvature generation during CME. For this purpose, ARPE cells overexpressing eGFP-labelled clathrin light chain were imaged by near simultaneous epifluorescence (Epi)-TIRF (total internal reflection fluorescence) microscopy [39] (Fig. 1B). Images acquired from the TIRF channel had higher signal-to-noise (SNR) ratio and were used for tracking clathrin intensities over time [13], whereas the Epi-channel more accurately reflects the rate of accumulation of clathrin at each CCP. *Bona fide* CCPs were identified from all the observed tracks using an unbiased filter of the intensity fluctuations that represent proper clathrin coat assembly [28]. Matching eGFP intensities were extracted from low SNR Epi images using primary/subordinate channel tracking [13]. The resulting CCP intensity traces from the two channels within a lifetime cohort of 24 *−* 40 seconds were aligned and averaged [40], and logarithmically transformed to give average traces of the invagination depth (Δ*z*) of the CCPs’ center-of-mass [39] (Fig. 1C-D). We chose the 24 *−* 40*s* cohort to reflect the invagination behavior of the most frequent tracks among *∼* 5 *×* 10^4^ CCPs over *>* 10 cells per each condition. See **Methods** for more details.

Clathrin assembly lymphoid myeloid leukemia protein (CALM), a phosphatidylinositol binding clathrin assembly protein, and epsin are pioneer proteins [1] that function at early stages of CME: both bind clathrin and are capable of bending membranes [41, 21]. We therefore examined the effect of siRNA-mediated knock-down (KD) of CALM and epsin1 (Fig. S1A-B) on Δ*z*. In negative control siRNA-treated cells Δ*z* ascended over time and reached *∼* 50*nm*, as expected for an average *∼* 100*nm* diameter CCV. In contrast, in CALM KD cells Δ*z* remained flat at *∼* 0*nm* (Fig. 1C). This KD effect is fully rescued by over-expressing CALM (Fig. S1C). KD of epsin1 also decreased initial curvature generation, albeit to a lesser extent (Fig. 1D). These defects in curvature formation indicate that CALM is required for and epsin1 contributes to the initiation of curvature at CCPs.

## A mechanochemical model for clathrin-coated pits

A computational model that incorporates the clathrin basket, the plasma membrane and EAPs’ curvature generation was developed for studying CME as a molecularly and mechanically integrated system. We used the preceding experimental findings in our model as a guide for setting the initial membrane morphology before simulating subsequent coat assembly and further membrane bending. To compute the evolution of membrane curvature during CME, we adopted a recent algorithm for simulating membrane morphological dynamics under external force based on triangular meshes [42] (Fig. 2A). The algorithm relies on a representation of the membrane as a mesh of discrete triangular units. As the membrane deforms, efficient remeshing is implemented to dynamically adjust the mesh configuration and to reflect membrane deformability without reducing mesh quality. This ensures accurate computing of the membrane’s bending energy throughout CCV formation [42]. Individual CME events are modeled as taking place on a complete sphere reflecting the closed topology of real cells (Fig. 2A). This avoids the complication of using truncated boundary conditions [43] and enables more accurate computation of bending forces [42]. CME events are initiated at the bottom of the sphere. To reduce computational load, the dimension of the model cell relative to a CCP was considerably smaller than for a real cell, but still large enough to enclose an entire CME event and to exclude non-CME related curvature effects. To set initial conditions for control cells, we introduced some initial curvature at the location of a future CME event prior to clathrin assembly. For the CALM KD cells, we introduced a flat region as the initial condition. For epsin1 KD cells, we introduced a degree of curvature between control and CALM as the initial condition (Fig. 2B). See **Methods** and (Fig. S2A) for more details and the units used in the model.

**Figure 2:**
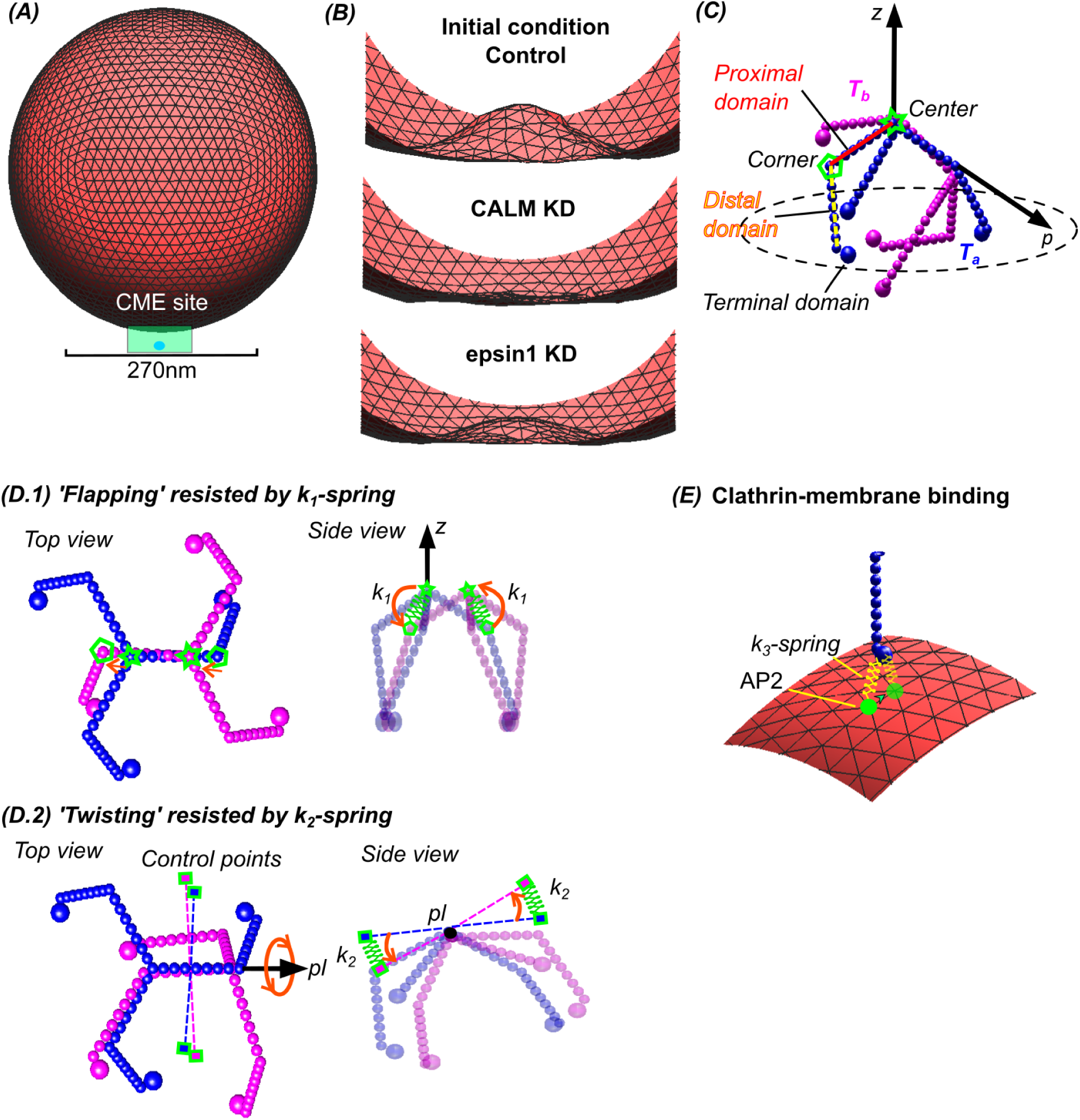
A mechanochemical model for clathrin-coated pits. (A) Boundary of simulated membrane and CME site. (B) Experimentally-guided initial membrane morphologies for control, CALM KD and epsin1 KD. (C) Coarse-grain model of clathrin triskelion composed of 52 spheres [45]. (D.1) Top and side view of ‘flapping’ flexibility and the springs connecting centers (star) and corners (pentagon) with coefficient *k*_1_ (green). (D.2) Top and side view of ‘twisting’ flexibility and the springs connecting control points outside triskelia with coefficient *k*_2_ (green). (E) Interaction between membrane and clathrin through AP2 (green) and connecting spring with coefficient *k*_3_ (yellow). Transparent AP2 and spring indicate a move of the clathrin terminal domain that reduces *k*_3_ spring energy.

### Flexibility of clathrin assembly

The experimentally observed requirements for CALM and epsin1 in initial curvature generation are consistent with previous findings in cells that clathrin-assembly alone was not sufficient to generate curvature [13], which in turn is consistent with the known flexibility of clathrin baskets [33, 44]. To reflect such flexibility and to compute the interactions between individual clathrin molecules, we adopted a coarse-grained approach that structurally describes the triskelion by chains of spheres [45] (Fig. 2C). One triskelion consists of a center sphere and three symmetrically attached arms, each consisting of one proximal, one distal and one terminal domain represented by 17 spheres in total. Multiple triskelia can self-assemble into polygonal baskets via interactions between their proximal domains that form lattice edges [44], a simplified but geometrically concrete representation of the high-resolution clathrin structure [46]. We described such binding using four harmonic springs to reflect the previously mentioned flexibility, a simpler version of the previously proposed schemes of clathrin flexibility [44, 47]. Specifically, we defined a spherical coordinate system where the center of a triskelion *T_a_*(blue in Fig. 2C) points in the +*z* direction, and the three terminal domains of *T_a_* remain on the equatorial plane. *T_a_* binds to another triskelion *T_b_* via interactions between their proximal domains. The first two springs, characterized by coefficient *k*_1_, connect the central sphere (marked with a star) of *T_a_*to the ‘corner’ sphere (marked with a pentagon) of *T_b_*. Conversely, they connect the center of *T_b_* to the corner of *T_a_* (Fig. 2D.1). The two *k*_1_ springs controlled the rotational motion of *T_b_* relative to *T_a_*about the mid-point between the two triskelia defining a ‘flapping’ flexibility.

The second two springs, characterized by coefficient *k*_2_, connect two imaginary control points outside *T_a_* to another two control points outside *T_b_*, acting as two weightless balancing poles (Fig. 2D.2). The relative positions from the control points to *T_a_* and *T_b_* were fixed so that the resulting spring forces and torques were entirely applied to the two triskelia. The two *k*_2_ springs mainly controlled the rotational motion of *T_b_* about the axis (*p*) defined by a proximal domain of *T_a_*, reflecting a ‘twisting’ flexibility [47]. All the springs in this work yield zero resting length, i.e. *l*_0_ = 0, and the total spring energy is equivalent to the chemical energy preserved in a clathrin basket. Note that both *k*_1_ and *k*_2_ springs penalize a third ‘sliding’ flexibility (Fig. S2B) that could lead to unrealistic polygonal structures with *»* 6 edges. Flat clathrin assemblies consist predominantly of hexagonal structures [20]: ‘sliding’ is thus implausible. On the other hand, ‘flapping’ flexibility is plausible, and can be manipulated by lowering *k*_1_ to flatten the hexagonal lattices. Increasing ‘flapping’ is independent on *k*_2_ springs (Fig. S2C); therefore, *k*_2_ is fixed at a high value to prohibit ‘sliding’ without affecting ‘flapping’. Next, we integrated the chemical energy from clathrin basket assembly with the mechanical energy from the bent membrane and computed the time evolution of the two components from initial configuration to steady state. The membrane and clathrin were mechanically connected via adaptor proteins, specifically AP2 [8], which are represented as particles occupying multiple vertices on the membrane. We used another spring with coefficient *k*_3_ to connect the AP2-occupied vertices and the terminal domains of the clathrin triskelia (Fig. 2E). This third spring reflects the effect of triskelia pulling or pushing the membrane. This tugging works against triskelion-triskelion binding. Thus, the *k*_3_ springs transfer energy from the clathrin coat assembly to the membrane and vice versa they impose mechanical constraints by the membrane on the assembly process. An AP2 particle moved from the membrane vertex it occupied only when moving to a neighboring vertex would reduce the *k*_3_ spring energy (green dots and yellow springs in Fig. 2E). This relocation reflected the sheer-stress-free property of the membrane.

Altogether, the described mechanochemical system represents individual clathrin triskelia as rigid bodies free to undergo translational Brownian motion determined by the spring forces, and rotational Brownian motion determined by the torques derived from those forces [48]. These include the *k*_1,2_ spring forces and torques from the clathrin-clathrin interaction, and the *k*_3_ spring force and torque from the clathrin-membrane interaction. On the other hand, individual vertices of the membrane, simply treated as particles, underwent translational Brownian motion. This motion is determined by the *k*_3_ spring force from the clathrin-membrane interaction, and the internal force from the membrane itself caused by bending (dominating term), lipid reorganization, osmotic pressure and tension [42]. See **Methods** for more details.

## Interplay between membrane curvature and basket structure

The membrane vertices dynamically depicted morphological changes upon interacting with a clathrin basket. Given the initial membrane morphology for a particular molecular configuration, we next let clathrin triskelia assemble one by one. Clathrin baskets were able to attach closely to both curved and flat membranes owing to the flexible interactions among triskelia and between triskelia and membrane. Most of the springs were not resting at their natural length *l*_0_ = 0, thus reserving some energy to deform the membrane. We stopped clathrin recruitment after 20 triskelia and relaxed the clathrin basket and the membrane together until equilibrium was reached. Separation of clathrin recruitment and relaxation was necessary to reduce computational time. Moreover, because considerable debate remains as to whether the clathrin basket assembles before or along with curvature formation [14], we chose to finish the assembly first then simulate the dynamics of the morphologies. See **Methods** for more details.

Modeling coat assembly at the three initial membrane morphologies reflecting control, CALM and epsin1 KD resulted in distinct pentagon-hexagon structures (Fig. 3A), which relaxed into different final morphologies (Fig. 3B). Emerging from this mechanochemical configuration, the 20 triskelia assembled into four pentagons in the control cells, but only one in the CALM KD cells, and three in the epsin1 KD cells (Fig. 3A, asterisks). As a result, the basket in the control cells eventually created sufficient curvature to successfully enclose a membrane vesicle (Fig. 3B), but the basket in the CALM KD cells stayed relatively open and flat, and the basket under epsin1 KD reached lower curvature compared to control. Importantly, these differences match our experimentally observed curvature defects of KD of curvature-generating pioneer EAPs, as well as the relative effects of CALM and epsin1 KD on CME and CCP dynamics [9, 28]. They also reveal positive feedback between initial membrane curvature and the incorporation of pentagons required for curvature-generation by nascent clathrin baskets.

**Figure 3:**
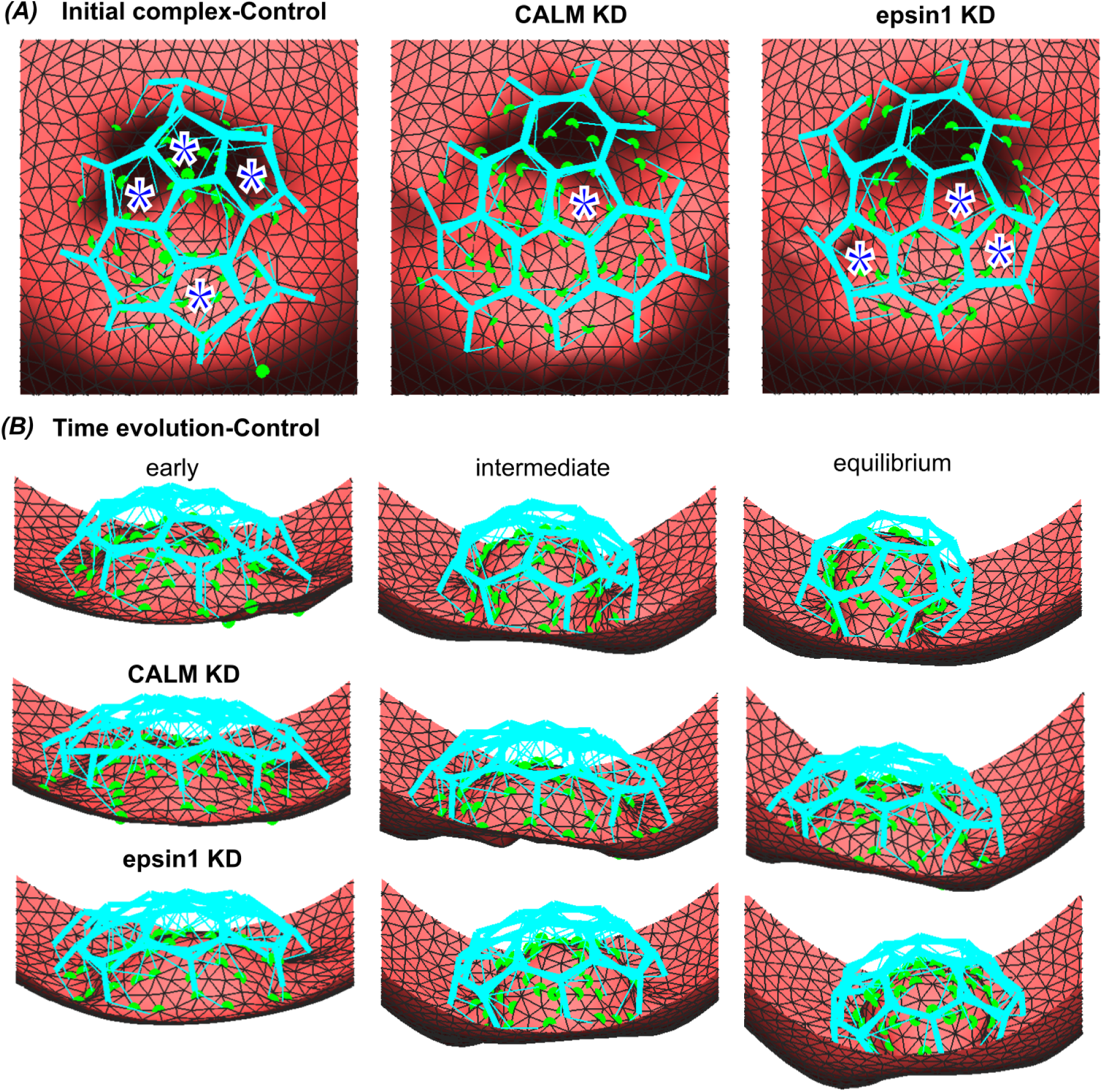
Interplay between membrane curvature and basket structure. (A) Initial complex simulated for clathrin and membrane of control, CALM KD, and epsin1 KD conditions. Asterisks indicate pentagons. (B) Evolution of the three complexes over time. The path to and the final equilibrium state are governed mechanochemical interaction of membrane and clathrin basket.

## CCPs evolve into two states—*Open* and *Closed*

We next exploited the model to investigate possible differences between abortive and productive fates of CCPs. For this purpose, we explored whether multiple equilibrium shapes of clathrin baskets exist and whether these shapes depend on the basket strength, which might vary under different molecular conditions (*i.e.* as the unimolecular clathrin ‘basket’ becomes a multi-component clathrin coat). The basket strength was varied computationally by tuning the spring coefficients. As discussed above, the *k*_2_ springs must be strong (*k*_2_ *»* 0) to avoid unreasonable clathrin structures; and the *k*_1_ springs could be lowered for generating flat lattices (see **Methods** and Fig. S2B-C). Therefore, we fixed *k*_2_ = 200 and treated *k*_1_ as a free parameter to adjust basket strengths. Additionally, we defined a dimensionless tightness factor

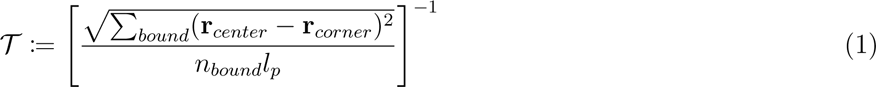

as the property to differentiate various equilibrium shapes of clathrin baskets. *T* monotonically reflects the tightness of the basket: the larger *T* is, the closer (*i.e.* tighter) clathrin triskelia bind to each other. *T* incorporates the summation of all the bound center-to-corner pairs, and is normalized by the number of pairs *n_bound_* and the constant length of the proximal domain *l_p_*. We also set *k*_3_ = 100 at a high value to let clathrin baskets effectively reshape the membrane.

Starting with an initial control membrane morphology, configurations with low *k*_1_ = 10 evolve to an *Open* basket equilibrium (Fig. 4A, Video 1). In contrast, at high *k*_1_ = 100, we observed *Closed* baskets at equilibrium (Fig. 4B, Video 2). To examine the molecular configuration of these states, we implemented a high-resolution, realistic visualization of the clathrin triskelia and lipid-based membrane following the physical scales indicated by the Protein Data Bank (rcsb.org) using a customized algorithm, see **Methods** for more details. Importantly, the outcome of these simulations in terms of basket shapes do not linearly depend on *k*_1_; that is, the baskets do not continuously loosen while decreasing *k*_1_. Instead, the curve *T* vs *k*_1_ displayed a sigmoidal behavior with a sudden transition from open to closed shapes as *k*_1_ crossed a critical value, *k_c_* (Fig. 4C). Interestingly, at *k*_1_ = *k_c_*, both *Open* and *Closed* states could be observed, indicating the existence of a switch-like behavior known to regulate biological systems at many scales, from neuronal synchronization to bird flocking [49]. Specifically, for a critical value *k_c_* = 30, CCPs coexist in an *Open* state (*T <* 20) or *Closed* state (*T >* 30). However, moving *k*_1_ beyond *k_c_* does not significantly change *T*, suggesting that a single closed state for these baskets exists. The two states obtained through time evolution are consistent with a reductionist model describing equilibrium membrane morphologies under molecularly induced mechanical manipulations [37]. Additionally, repeating simulations at *k*_1_ = *k_c_*, yielded a bimodal distribution for *T* (Fig. 4D), in agreement with recent super-resolution microscopy images of static CCPs [50].

**Figure 4:**
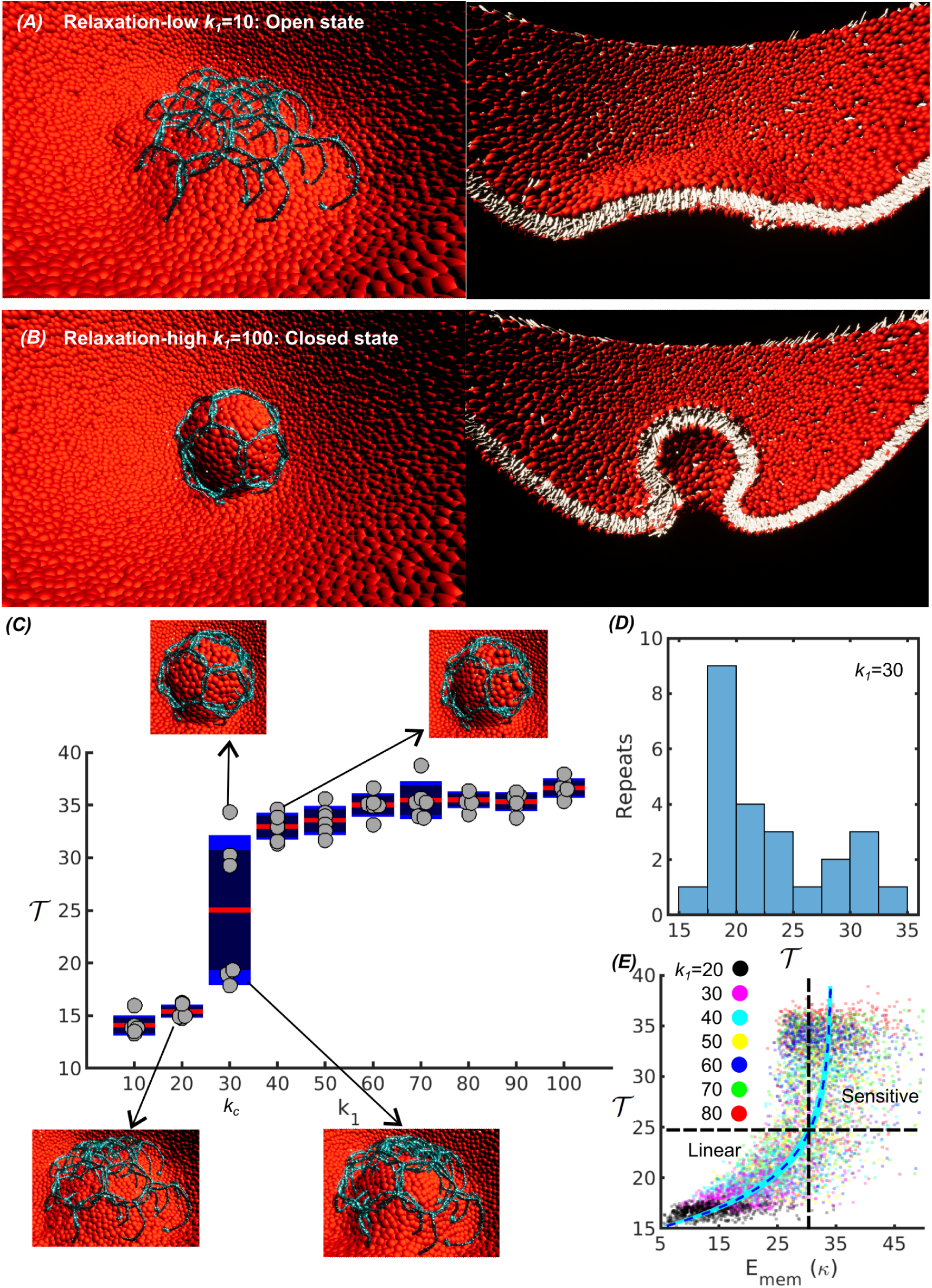
CCPs evolve into two states—*Open* and *Closed*. (A,B) Realistic high resolution visualization of clathrin-membrane complex equilibrated (A) at *k*_1_ = 10 (*Open* state) and (B) at *k*_1_ = 100 (*Closed* state). Left column: top view; right column: side view. (C) Clathrin basket tightness, *T* (Eq. 1), plotted against various values of *k*_1_. The curve shows a sharp transition between *Open* and *Closed* state at *k*_1_ = *k_c_* (six simulation repeats per *k*_1_ value). (D) Bimodal histogram of *T* for simulations at *k*_1_ = *k_c_* (*k*_1_ = 30). The stochasticity of of outcomes originates in randomized remeshing decisions of the membrane representation during the simulation. (E) Total membrane bending energy at the vertices covered by clathrin *T* vs *E_mem_* as solid dots in units of the membrane bending modulus *κ*. Different colors represent data collected from time evolutions at various *k*_1_ values. Dash line (blue) and band (cyan) represent an exponential fitting and its 95% confidence.

The two states emerging at *k_c_* are due to the membrane’s sensitive response to basket closure—the bending energy (*E_mem_*) at the membrane vertices near the clathrin baskets are nonlinearly related to the basket tightness (*T*). To reveal this relation, *>* 300 values of *E_mem_*with *T* were chosen to fully represent the time evolution of a given CCP from its initial to equilibrium state. We examined multiple *k*_1_ = 20, 30*, …*70 values (six evolutions per each *k*_1_) to incorporate both *Open* and *Closed* scenarios. Each data point of (*E_mem_*,*T*) is represented as a dot colored according to *k*_1_ in Fig. 4E. After pooling all the data points, we show that when *E_mem_*is small, increasing *E_mem_* corresponds to a nearly linear increase of *T*, indicating a more energetically demanding stage of closure. However, when *E_mem_* 2: 30*κ* (where the bending modulus of the membrane *κ* = 100*k_B_T*), small increases in *E_mem_* correspond to a dramatic increase of *T*, indicating a nearly effortless closure of baskets at *k*_1_ *≥ k_c_*. Baskets with *k*_1_ *< k_c_* are gated by the energetic threshold. The sensitive relation between the membrane and the baskets governs the bi-state transition around *k_c_*.

## The endocytic checkpoint is a mechanochemical switch

Biologically, the *Open*/*Closed* switch delineates the state boundary between abortive and productive CCPs. Based on our model, we speculate that cells operate with clathrin coats assembled near the critical value *k*_1_ *∼ k_c_*. Small changes in *k*_1_ can thus control the abortive vs productive fate of CCPs. We propose that the exquisite sensitivity of the switch defines the mechanism of the endocytic checkpoint. Transitions from *Open* to *Closed* can occur with the recruitment of clathrin or AP2-binding EAPs that strengthen the clathrin basket to alter *k*_1_ [11]. Additionally, EAPs that strengthen interactions between clathrin baskets and the membrane, e.g. by binding membranes, AP2 and/or clathrincan affect *k*_3_ springs and impact the relation *k*_1_ vs *T* . Cutting *k*_3_ in half leads to lower curvatures because the increased compliance of the springs allows the membrane to\ depart from the basket geometry and relax in a flatter configuration with smaller bending force. As a result, open baskets at *k*_1_ = 30 and even some at *k*_1_ = 20 become closed (Fig. S3A-B). Conversely, doubling *k*_3_ raises geometric constraints imposed by the basket onto the membrane, thus causing larger curvatures and larger bending forces that prevent open *k*_1_ = 30 baskets to close (Fig. S3C-D). Lastly, EAPs that induce membrane curvature could alter *k_c_* by sharing some energetic burden with clathrin and allowing baskets with *k*_1_ *< k_c_* to close. This could explain the observed relationship between membrane curvature and clathrin assembly [10, 11, 12].

Following these findings, we therefore computationally evaluated whether the fate-determining switch that emerges from the competition between basket closure and membrane resistance against bending could be influenced by EAPs that induce curvature to assist clathrin. For this, we introduced *k*_4_ springs (Fig. 5A) to parameterize additional curvature generation at the clathrin lattice. Curvature generation by EAPs would force compaction of the internal membrane meshwork. Thus, these *k*_4_ springs pull neighboring AP2-occupied membrane vertices towards a controlling center that extends extracellularly by a certain distance *r_w_*. Indeed, these added forces help the clathrin baskets to bend the membrane. Under weak *k*_4_ springs (*k*_4_ = 10), *T* values at *k*_1_ *≤ k_c_* were already significantly increased. Consequently, some originally *Open* baskets with *k*_1_ *< k_c_*(*k*_1_ = 20, i.e. modeling increased EAP-imposed curvature) and all of the open baskets with *k*_1_ = *k_c_* = 30 became closed (Fig. 5B.1, S4). The baskets with *k*_1_ *k_c_* (*k*_1_ = 10) stayed open. Under stronger *k*_4_ springs (*k*_4_ = 20), all of the *k*_1_ *< k_c_* (*k*_1_ = 20) baskets and even a *k*_1_ *k_c_* (*k*_1_ = 10) basket closed (Fig. 5B.2, S4). Importantly, the observed switch-like behavior, and the co-existence of two states, persist independent of *k*_4_. Thus, curvature generating EAPs could influence the switch-like endocytic checkpoint by increasing *T* at *k*_1_ *< k_c_*, or, effectively, by lowering *k_c_*, to promote basket closure and thus productive CCPs.

**Figure 5:**
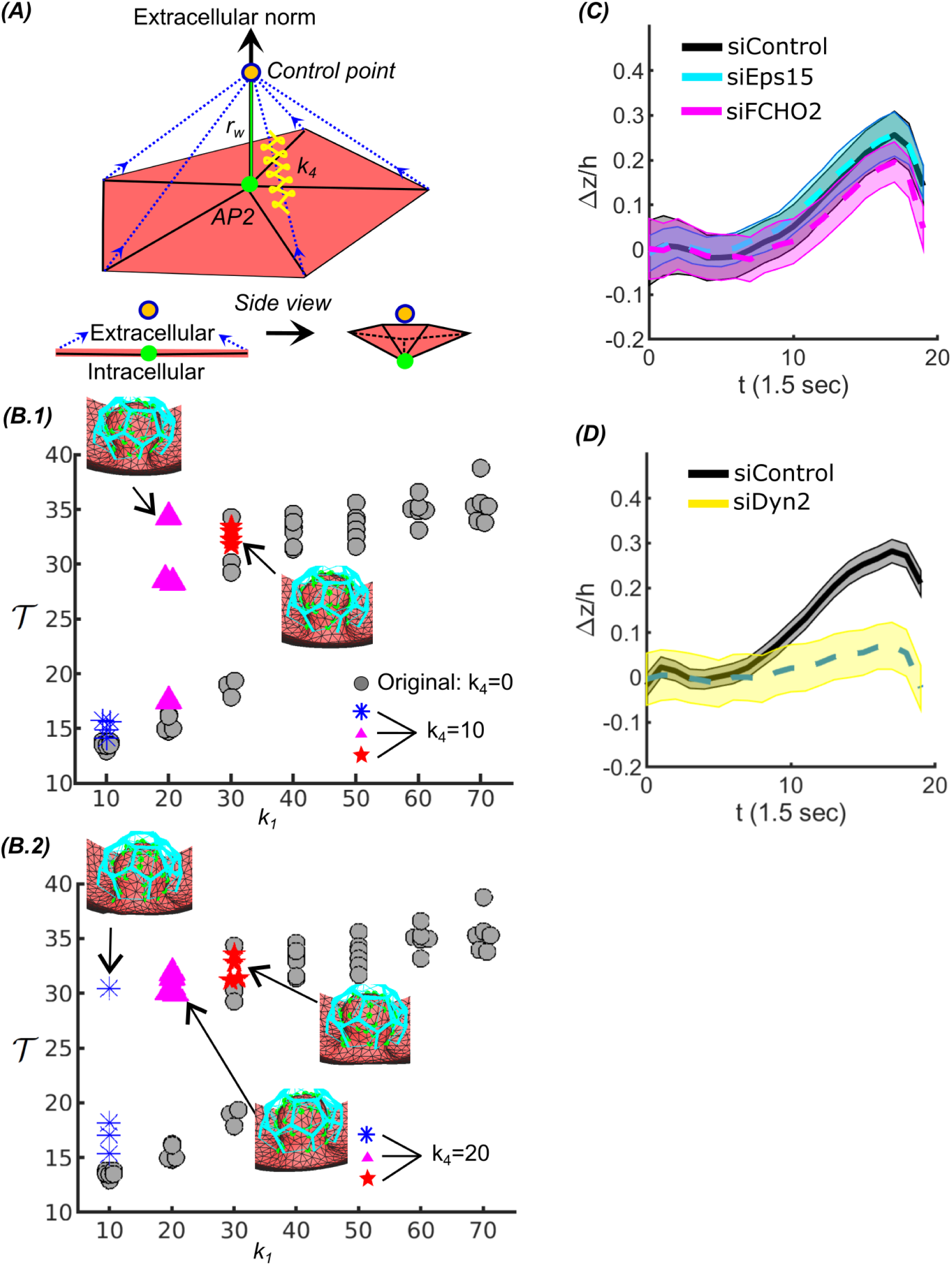
The endocytic checkpoint is the emergent property of a dynamic molecular system describing the mechanochemical interaction between membrane and clathrin basket. (A) Clathrin basket assembly is aided by EAP-dependent curvature generation. *k*_4_ springs mimic EAP- mediated curvature generation at edges of clathrin baskets. (B) *k*_1_ vs *T* curve shifted near *k_c_* when adding curvature by *k*_4_ springs with (B.1) *k*_4_ = 10 and (B.2) *k*_4_ = 20. (C) Normalized and averaged invagination depth, Δ*z*(*t*)*/h*, measured in siControl (pooled from *n* = 6.3 *×* 10^3^ traces from *m* = 9 cells), Eps15 KD (pooled from *n* = 6.3 *×* 10^3^ traces from *m* = 10 cells) and FCHO2 KD (pooled from *n* = 4.1 *×* 10^3^ traces from *m* = 10 cells) conditions. (D) Normalized and averaged invagination depth, Δ*z*(*t*), measured in siControl (pooled from *n* = 11.8 *×* 10^3^ traces from *m* = 15 cells) and Dynamin2 (Dyn2) KD (pooled from *n* = 4.7 *×* 10^3^ traces from *m* = 15 cells) conditions. Solid/dashed lines indicate the average traces, shaded bands indicate 95% confidence intervals.

*A role for dynamin2 in the endocytic checkpoint*. Dynamin is an EAP best known for its roles during membrane fission [51]; however numerous studies have implicated an earlier function for dynamin2 (Dyn2) in regulating the endocytic checkpoint [4, 13, 2]. In experiments with endogenously labelled Dyn2, both Dyn2-positive and -negative CCPs were observed [13] and statistical analyses of thousands of pits revealed that Dyn2-negative CCPs are predominantly abortive. In line with this data, siRNA-mediated knockdown of Dyn2 resulted in a decrease in stabilized CCPs and an increase in short-lived abortive CCPs [52]. In further support of its early role in CME, Dyn2 is present on nascent and flat CCPs [53, 32] and high-resolution correlative light and electron microscopy has positioned Dyn2 towards the edges of flat and domed clathrin lattices, along with FCHO and Eps15 [32]. Finally, results from a comprehensive CRISPRi-mediated knockdown screen of 67 EAPs, revealed common phenotypes between Dyn2 and several pioneer EAPs (e.g. CALM. Eps15, epsin1, FCHO1, NECAP2), namely a lower initiation rate and smaller fraction of stable CCPs [9].

Based on their spatial and phenotypic similarities, we tested the effects of siRNA-mediated KD of Dyn2, Eps15 and FCHO2 (Fig. S5A-C) on CCP invagination depth (Δ*z*) using our Epi-TIRF assay. Unexpectedly, KD of the known curvature-generating protein FCHO2 did not significantly inhibit CCP invagination, nor did KD of Eps15 (Fig. 5C). In contrast, siRNA-mediated KD of Dyn2 resulted in a strong reduction in CCP invagination depth (Δ*z ∼* 0) similar to CALM KD (Fig. 5D). This effect was fully rescued by re-introduction of Dyn2 (Fig. S5D). The observed Dyn2-requirement for curvature generation earlier than its membrane-fission function [54] could result either from known direct (through its wedging function [55]) or indirect (through its interactions with curvature generating EAPs [7]) activities of Dyn2. These results establish that Dyn2 plays a key role in regulating the endocytic checkpoint by relieving the constraint on basket strength to allow baskets with *k*_1_ *≤ k_c_* to close (Fig. 5B).

## Conclusion

Our combined experimental and computational findings define the long-proposed concept of an endocytic checkpoint as an emergent property of the mechanochemical interaction between membrane and clathrin machinery. Using a realistic model of these interactions adjusted to experiments, we identify two states of CCPs according to the shape of clathrin baskets: an *Open* state for weak baskets and a *Closed* state for strengthened baskets. These two states match our previous perception of the abortive and productive fates developed based on the statistics of CCP lifetime. We speculate that plasma membranes and the clathrin baskets that assemble upon them are poised at *k*_1_ *∼ k_c_*, hence allowing for the regulation of CME by EAPs that affect membrane curvature and/or the strength of the assembled basket.

The presented data and accompanying model jointly resolve three long-standing puzzles in the field of CME: i) We find that nascent CCP progression is gated by an emergent mechanochemical property of the energy balance between clathrin basket assembly and membrane bending. Thus the elucive endocytic checkpoint is not a molecular machine, but a biophysical characteristic of CCP maturation; ii) The sensitive switch-like behavior of the endocytic checkpoint offers an explanation for how functionally diverse EAPs can determine the fate of abortive or productive CCPs; iii) We discover Dyn2 as a key regulator of early curvature generation and provide a mechanism for its observed ability to regulate the endocytic checkpoint. Together these discoveries explain the resilience and observed compensatory behavior of CME [13, 28, 14] in that the mechanochemical switch-like behavior of the checkpoint is exquisitely sensitive to the interplay between membrane curvature and the strength of the clathrin basket, either of which can be modulated by functionally diverse EAPs.

## Methods

### Cell lines

ARPE19-HPV16 cells were obtained from ATCC and cultured in Gibco DMEM/F12 medium (ThermoFisher, 11330032). A version of the cell line stably expressing eGFP-CLCa was generated as described in our previous work [56].

### siRNA transfection and rescue

Cells for siRNA interference were seeded on a six-well plate (250,000 cells/well) and transfected with siRNA after 6 hours. Two rounds of transfection were conducted over 3 days to achieve *>* 90% target protein knockdown on day 4 of the experiments. For each transfection, 100*µl* Opti-MEM with 110 pmol siRNA (tube 1) and 100*µl* Opti-MEM (Gibco) with 6.5*µl* Lipofectamine RNAi-MAX (Invitrogen) (tube 2) were incubated at room temperature for 5 min. Then, the reagents in tube 1 and 2 were mixed and incubated at room temperature for 15 min before adding them drop by drop to the cells. siControl: 5’-UUC UCC GAA CGU GUC ACG U -3’; siCALM: 5’-ACAGUUGGCAGACAGUUUA-3’; siDyn2 (3’UTR): a mixture of #1 5’-CUGCACUCUGUAUUCUAUUAAUAAA-3’, and #2 5’-GGUAUAUCAACUUCCCAUUAGCAGG-3’; siEPS15: 5’-AAACGGAGCUACAGAUUAUUU-3’; siFCHO2: 5’-CCAAGUGUGUAGAACAGGAGCGUUU-3’; si-epsin1: mixture of #1 5’-GAUCAAGGUUCGAGAGGCC-3’, #2 5’-GGAAGACGCCGGAGUCAUU-3’, #3 5’-GAACGUGCGUGAGAAAGCU-3’, and #4 5’-ACUAAUCCCUUCCUCCUAU-3’. CALM (Protein-Tech, 28554-1-AP), Dyn2 (SantaCruz, sc-166669), EPS15 (abcam, ab174291), epsin1(ZEN-BIO, R24229), FCHO2 (Genetex, GTX120444), and GADPH (ABclonal, AC033) antibodies were used for Western Blotting to confirm siRNA knockdown efficiency.

For CALM rescue experiments, we expressed wildtype CALM (with a myc-tag and siRNA-resistant coding sequence) using a retroviral vector (giftfrom David Owen) [41]. For Dyn2 rescue experiments, wildtype DNM2 with a N-terminal HA-tag was cloned into pLVx-IRES-puro vector with a distinct 3’ UTR. Corresponding retro- and lenti-viruses were produced in 293T cells and used to infect ARPE19-HPV16 eGFP-CLCa cells to achieve stable expression of exogenous CALM and Dyn2.

### Epi-TIRF microscopy imaging and quantification

Cells were grown on a gelatin-coated 35mm glass bottom ibidi dish (ibidi, #81218) overnight. Imaging was conducted with a 100*×* oil, NA 1.49 Apo TIRF objective (Nikon) mounted on an Eclipse Ti2 inverted microscope equipped with 1) a H-TIRF module for TIRF acquisition (penetration depth = 80*nm*), 2) a M-TIRF module for epifluorescence acquisition, and 3) an Okolab cage incubator to sustain 5% CO2 and 37*^◦^C*. For time-lapsed Epi/TIRF imaging, TIRF and epifluorescence signals were acquired nearly simultaneously at the frame rate of 0.66*frame/s*. The Nikon Perfect Focus System (PFS) was applied during imaging.

After imaging, *cmeAnalysis* [13] was applied for tracking fluorescently labelled particles, including gap closing [57]. Tracks that overlap with others or deviate from the properties of a diffraction-limited particle were excluded from the analysis. Tracks passing this filtering scheme were further classified into *bona fide* CCPs vs clathrin-coats by *DASC* [28], a fluctuation-based unbiased method. The tracks classified as CCPs were used for Δ*z* measurement. All image analysis software and parameter settings for Epi-TIRF analysis are available at https://github.com/DanuserLab/cmeAnalysis.

### Modeling

#### Membrane initiation

For simulations of membrane bending under the mechanical constraints imposed by the clathrin-coat we leveraged the recently published triangular-mesh-based method [42]. This method optimally balances the physical and geometrical conditions that govern the computing accuracy and efficiency. The dynamics of CCP morphology is captured by an algorithm that controls the mesh-based representation of the membrane based on physical potentials. Remeshing operations are initiated when either the triangular geometry or the local strain in the mesh degenerate, thus maintaining high mesh quality for accurately computing membrane-related interactions. To start a simulation a spherical membrane is slightly pulled by external control points that initiate curvature. Dependent on the molecular conditions (WT, CALM KD and epsin1 KD) the points were placed at different heights to generate different initial equilibrium geometries for further computation (Fig. 1F, Fig. S3A); see the method ‘*ClathrinMediatedEndocytosisInit* ’ in the class ‘@BiophysicsApp’ on GitHub: https://github.com/DanuserLab/biophysicsModels for further documentation. Throughout the paper the energy unit 1*k_B_T* and length unit 10*nm* are used for scaling.

#### Forces, torques, noises and dynamics

All spring forces were computed as **F***_s_*_12_ = *−k*(**r**_2_ *−* **r**_1_), indicating the force exerted in point 1 by an elastic connector to point 2. Other forces were numerically derived from the energy potential as **F** = *−*[*E*(**r** + Δ**r**) *− E*(**r**)]*/*Δ**r** (see Table S1). These potentials include volume and surface area constraints on the membrane, reflecting osmotic pressure and membrane tension; the Helfrich bending energy [58], reflecting the membrane bending force; and the internal energy of the membrane, reflecting the resistance to deformation. The latter also controls the mesh geometry, as described above. Implementing these forces requires an adaptive time step for enhancing computational efficiency. Detailed implementations and instructions of these forces, the remeshing and adaptive time step can be found in [42] and our GitHub repository. We treated clathrin triskelia as rigid bodies following [48]. Each triskelion consists of 52 spheres according to [45]. A Cartesian rotation vector **a** defined as the rotation coordinates [59] was implemented to describe rotational movement of the triskelia. As a result, the torque within the triskelia is derived from the rotational variation to the total spring energy 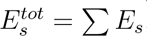, summing over all the pairs of connected points (see Table S1):

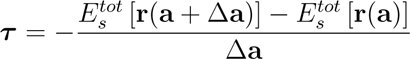

Here **r** indicates the coordinates of the points in the triskelia and **a** indicates the triskelia’s orientations in space [48].

All forces on the membrane vertices were included in a Langevin equation with adaptive time steps that efficiently describes overdamped Brownian motion, see [42]. The random force was eliminated to reduce the noise that affects the morphologies. All forces and torques on the clathrin triskelia were included in a rotational Langevin equation [48] with the random force eliminated. Also, the *k*_3_ spring force, and the forces derived from osmotic pressure and tension in a particular vertex were spatially smoothed by averaging over the immediately neighboring vertices. Removing the random forces and the spatial averaging reduced the impact of noise and numerical distortions associated with the discrete meshing, resulting in sharper transitions between the *Open* and *Closed* states. However, the remeshing process still implies on some random decisions during triangulation [42]. Therefore, for any given parameter setting multiple simulations were run to obtain statistically meaningful results. The dynamics of CCP formation is implemented in the application ‘*ClathrinMediatedEndocytosis*’.

For simulating a molecular wedge, *k*_4_ springs generate forces in the vertices neighboring an AP2-occupied vertex, pointing towards the control points above the vertex (Fig. 5A). These forces are balanced in the control point by a force exerted on the AP2-occupied vertex. For asymmetric mesh configurations, these two sets of forces generate also a torque about the connector between vertex and control point. However, for computational simplification we neglect this torque because of the usually nearly equidistant arrangement of the vertices pulled by *k*_4_ springs about the AP2 vertex.

#### Clathrin assembly

After initializing the membrane morphology, clathrin triskelia were placed on the membrane one by one. After a triskelion attaches to the membrane it was subjected to mechanical relaxation before the next, adjacent triskelion was placed. The newly added triskelion was relaxed with at least one proximal leg bound to the previous ones. During relaxation, the membrane was fixed. However, the feet of the triskelia were allowed to move to connect to membrane vertices under minimization of of the *k*_3_ spring energy. We limited the basket sizes to 20 triskelia for computational efficiency. Moreover, because of the size-invariance of the mebrane bending energy [58], the default basket of 20 triskelia bears a higher energy density than baskets with a larger size. Therefore, smaller baskets are more relevant to explore the functions of EAPs in modulating the checkpoint.

#### High-resolution visualization of simulation data in 3D

The 3D visualizations of clathrin-membrane interactions were generated using OmVisim, a 3D visualization software created by Omphalos Lifescience Inc. Each spatial-related property from the original Matlab simulation data was linearly transformed to correspond to the coordinate space of OmVisim, without altering the original shape and relative positioning of the objects. For clathrin visualization, a set of positions and orientations of clathrin was used to spawn a single triskelion mesh (PDB ID: 3IYV on Protein Data Bank; processed using PyMOL —https://pymol.org/2/). For membrane visualization, the triangle vertices of the membrane and interpolated positions between the vertices were used to approximate a membrane surface. Assuming a lipid bilayer depth of 7*nm*, another membrane surface was generated solely for visualization purposes. Then, lipid meshes (PDB ID: 8ND; processed using PyMOL and Blender—https://www.blender.org/) were spawned to populate the surfaces, resulting in a lipid bilayer visualization. To view cross-sections of the visualizations, a “null space” was introduced into the 3D environment where nothing is rendered. Visual parameters that are not derived from any simulation data, including lipid size and other stylistic properties like color, lighting, and camera positioning, were fine-tuned to enhance visual clarity.

## Acknowledgment

We thank Jenny (Qiongjing) Zou for software management, Madhura Bhave for data sharing and Milo Lin for helpful discussion. This work was supported by NIH grants (R35GM135428 to G.D. and R01GM73165 to M.M.) and the National Natural Science Foundation of China grant (No. 32200564 to Z.C.).

---

**Figure 6:** Video 1: scale-free animation of an open clathrin basket with interacting membrane.

**Figure 7:** Video 2: scale-free animation of a closing clathrin basket with interacting membrane.

**Table S1:**
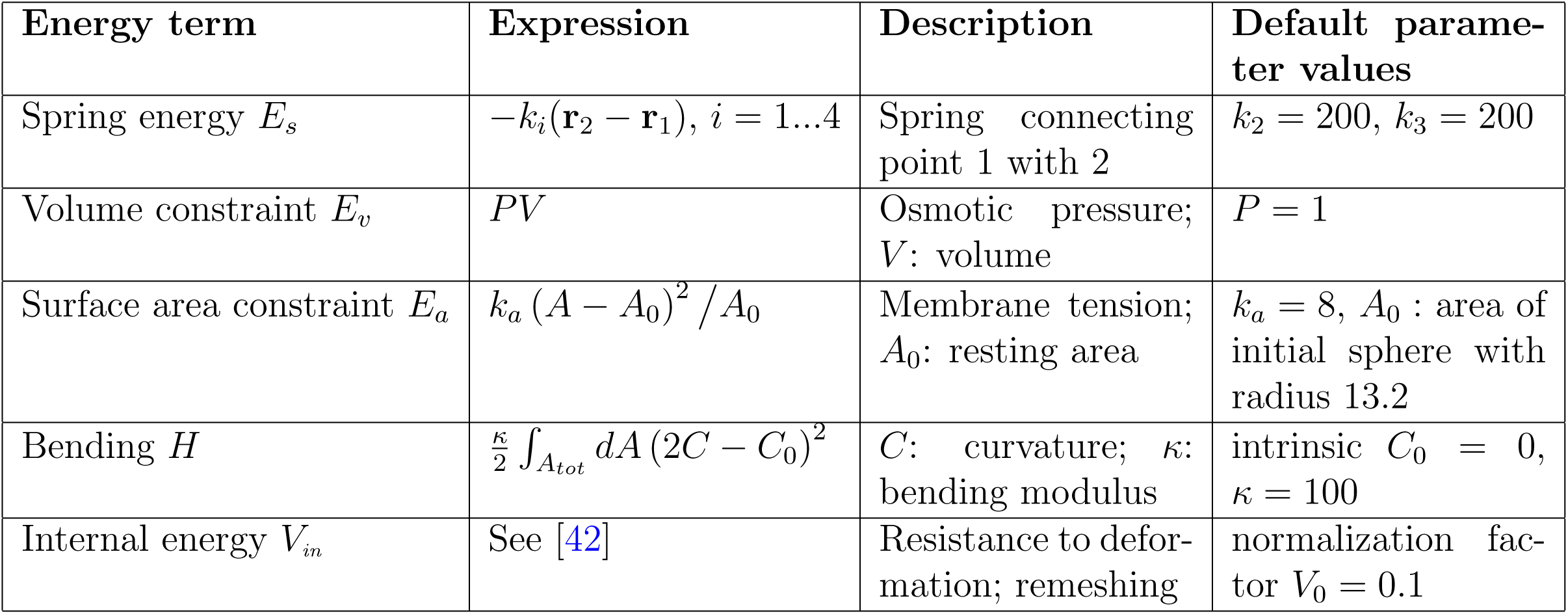
Energy terms and descriptions.

**Figure S1:**
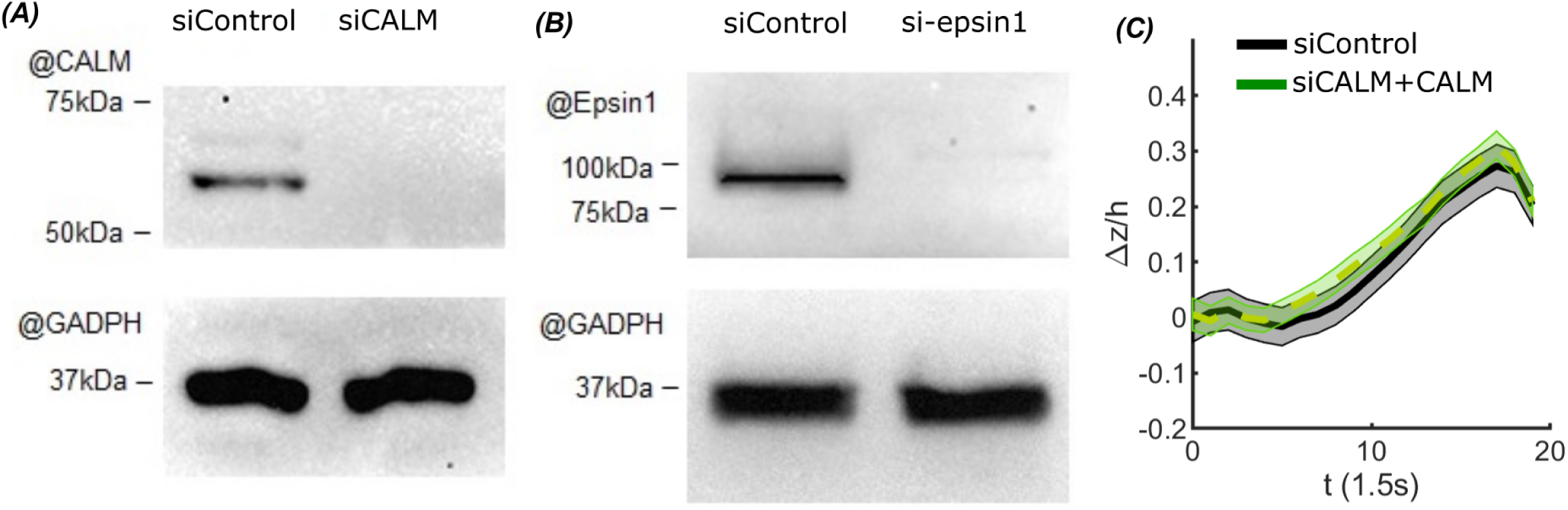
Knockdown efficiency of CALM and epsin1 and rescue effect of CALM. (A-B) Western blots of showing efficient KD of (A) CALM and (B) epsin1. (C) Depth of invagination (Δ*z*) phenotypes can be rescued by over-expressing exogenous CALM. CALM+: cells stably express exogenous CALM via lentiviral infection and siRNA knockdown of endogenous CALM. (C) Normalized and averaged invagination depth, Δ*z/h*, in cells overexpressing an siRNA-resilient form of CALM (siCALM + CALM; pooled from *n* = 14.0 *×* 10^3^ traces from *m* = 15 cells). For comparison, siControl in Fig. 1C is replotted.

**Figure S2:**
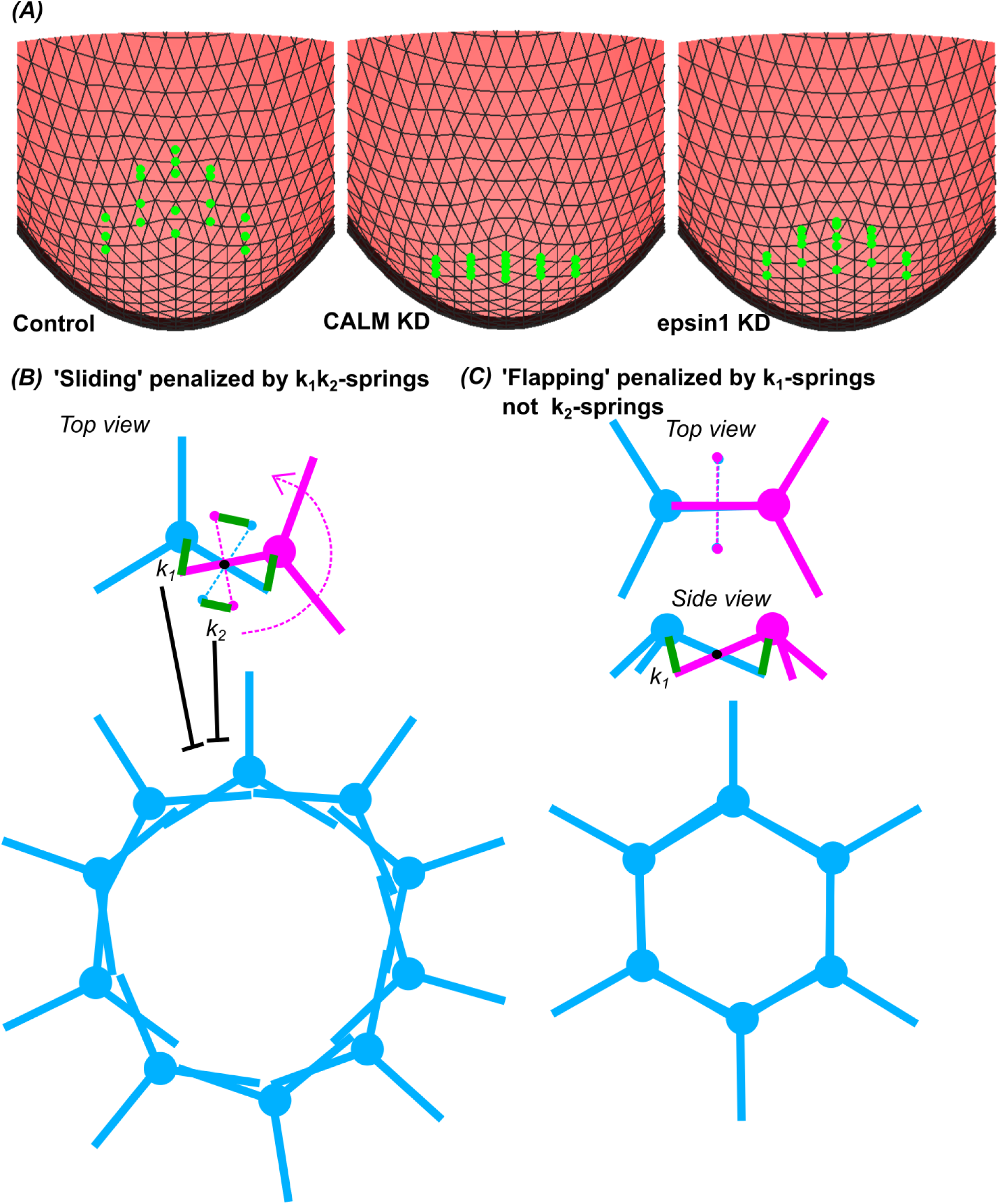
Initial morphology generation and clathrin’s flexibility and lattice structures. (A) Generation of initial morphologies by control points (green dots) pulling membrane vertices in control, CALM KD and epsin1 KD conditions. (B) ‘Sliding’ flexibility and implausible lattice structure. (C) ‘Flapping’ flexibility and plausible lattice structure.

**Figure S3:**
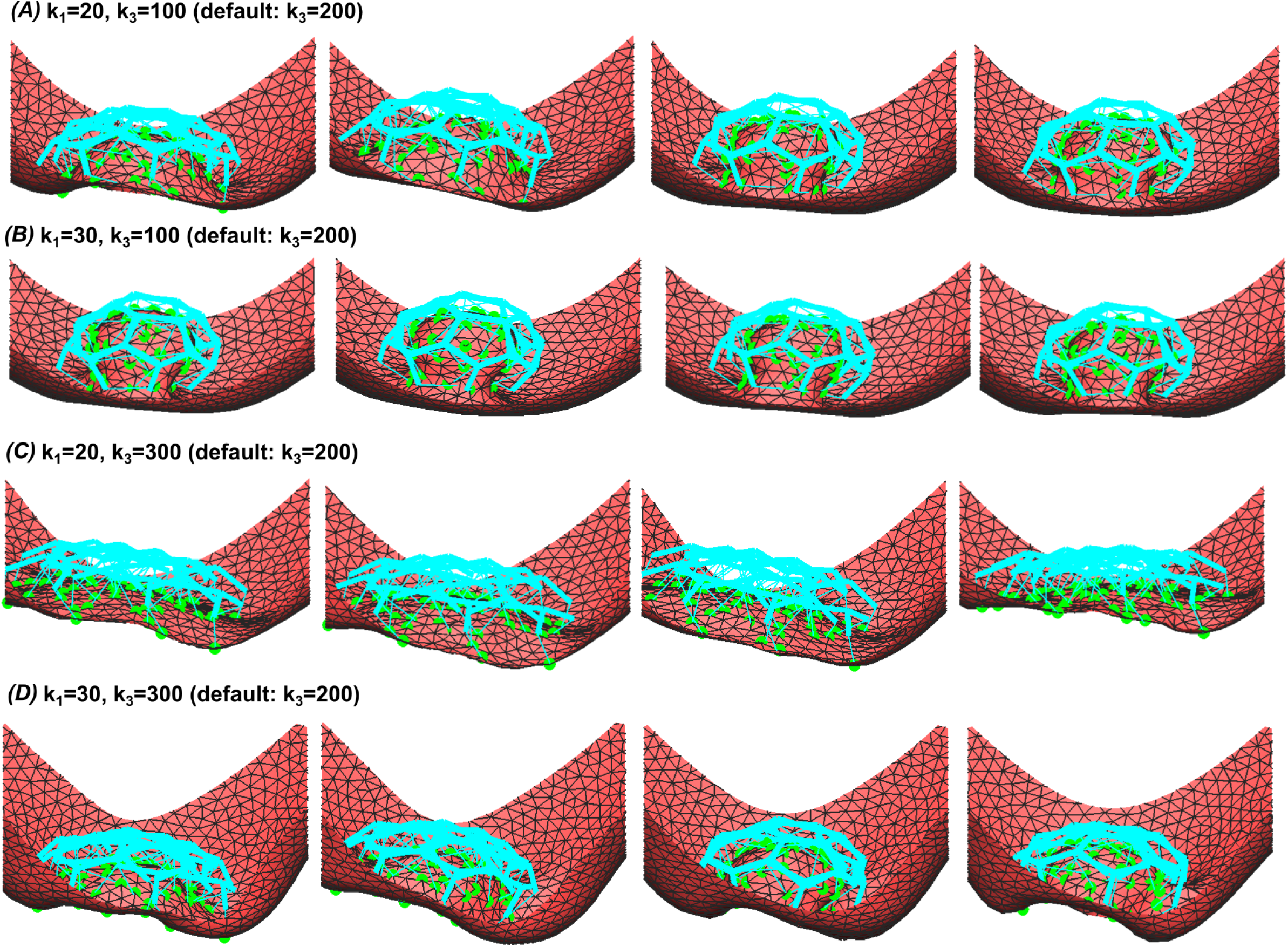
Equilibrium states affected by varied *k*_3_ springs. (A,B) Reduced *k*_3_ (*k*_3_ = 100) causes some originally *Open* baskets with *k*_1_ *< k_c_* to close (A) and all the originally *Open* baskets with *k*_1_ = *k_c_* close (B). (C,D) Increased *k*_3_ (*k*_3_ = 300) does not affect originally *Open* baskets with *k*_1_ *< k_c_* (C), but *Open* baskets with *k*_1_ = *k_c_* cannot close (D).

**Figure S4:**
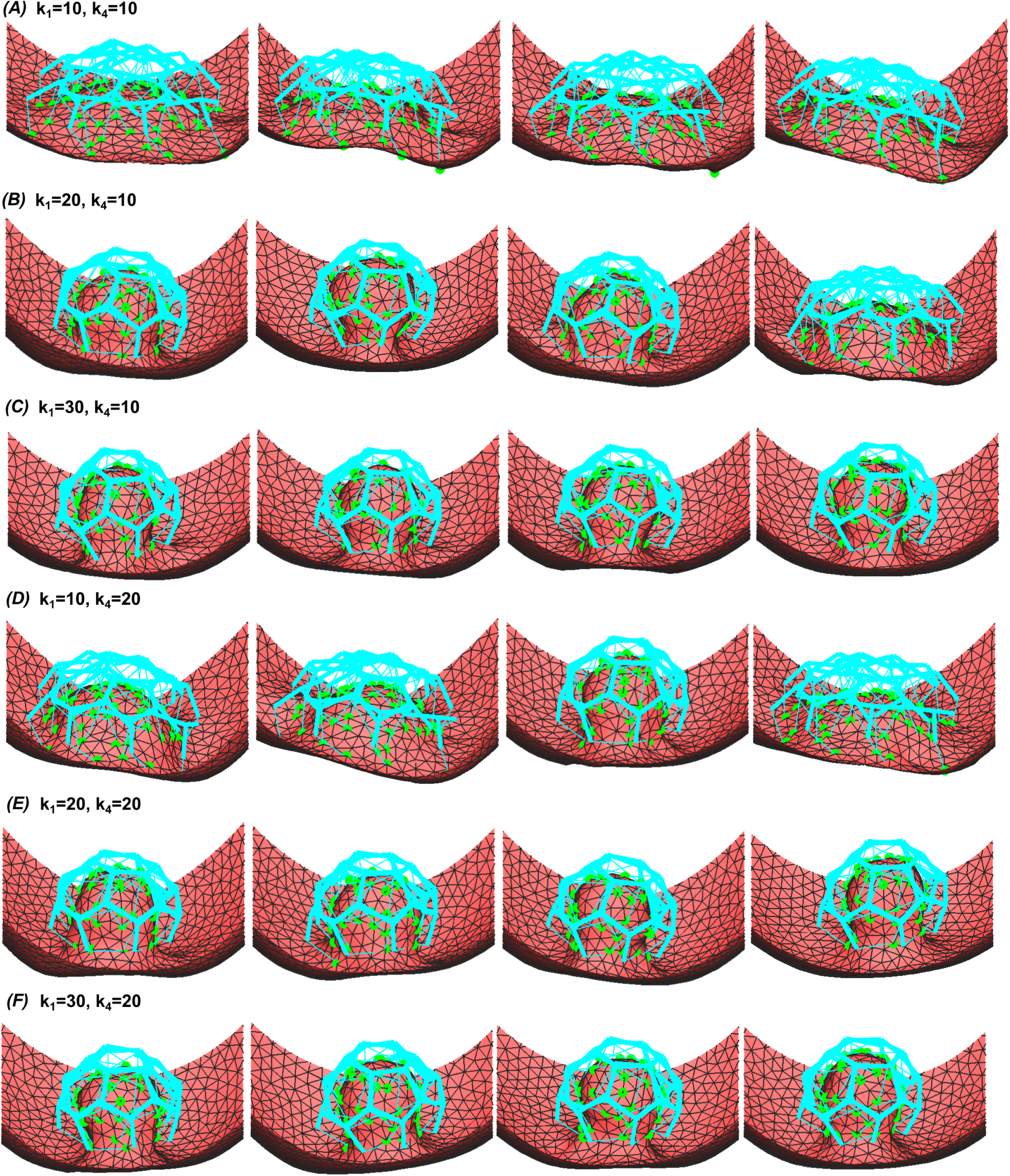
Equilibrium states affected by *k*_4_ springs. (A-C) Weak *k*_4_ springs (*k*_4_ = 10) (A) do not affect baskets with *k*_1_ = 10 *« k_c_*, but (B) cause some *k*_1_ = 20 *< k_c_*baskets to close, and (C) all of *k*_1_ = 30 = *k_c_* baskets to close. (D-F) Strong *k*_4_ springs (*k*_4_ = 20) (D) cause one *k*_1_ = 10 *« k_c_*to close, (E) most *k*_1_ = 20 *< k_c_* baskets and (F) all of *k*_1_ = 30 = *k_c_*baskets to close.

**Figure S5:**
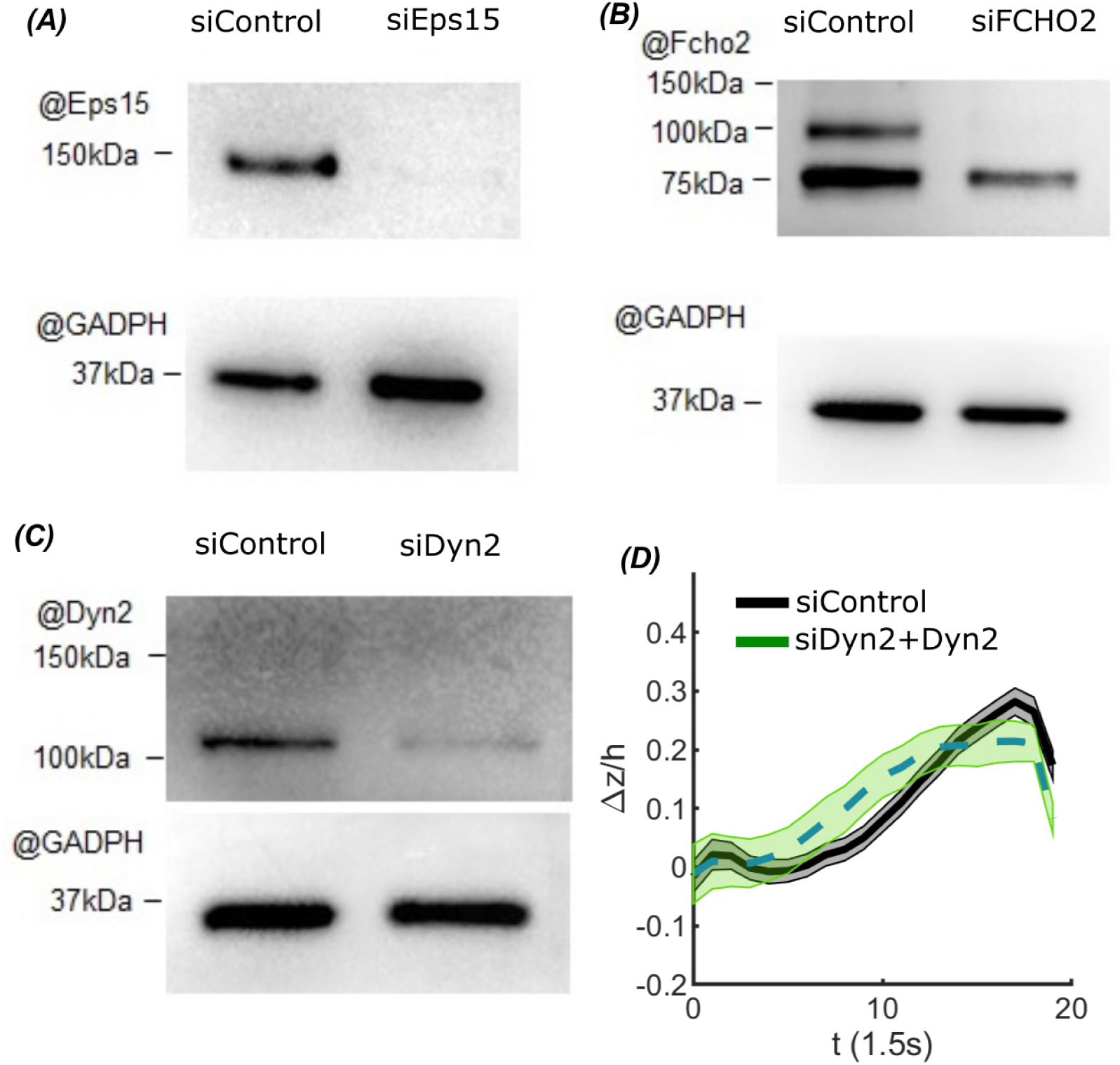
Knockdown efficiency of Eps15, FCHO2 and Dynamin2 (Dyn2) and rescue effect of Dyn2. (A-C) Western blots are showing efficient KD of (A) Eps15, (B) FCHO2 and (C) Dynamin2. (D) Normalized and averaged invagination depth, Δ*z/h*, in cells overexpressing an siRNA-resilient form of Dyn2 (Dyn2+; pooled from *n* = 15.7 *×* 10^3^ traces from *m* = 15 cells). Specifically, Dyn2+ cells stably express exogenous Dyn2 via lentiviral induction and 3’ UTR siRNA-mediated knockdown of endogenous Dyn2. For comparison, normalized and averaged invagination depth for control cell expressing a scrambled siRNA-vector are shown (pooled from *n* = 5.8 *×* 10^3^ traces from *m* = 11 cells).

## Notes

### Competing Interest Statement

The authors have declared no competing interest.

